# Resistance to clothianidin reduces the efficacy of SumiShield® 50WG, a neonicotinoid formulation for indoor residual spraying, against *Anopheles gambiae*

**DOI:** 10.1101/2020.08.06.239509

**Authors:** Caroline Fouet, Fred A. Ashu, Marilene M. Ambadiang, Williams Tchapga, Charles S. Wondji, Colince Kamdem

## Abstract

Chronic exposure of mosquito larvae to pesticide residues in agricultural areas is often associated with evolution of resistance to insecticides used for vector control. This presents a concern for the efficacy of clothianidin, an agricultural neonicotinoid qualified for Indoor Residual Spraying (IRS). Using standard bioassays, we tested if reduced susceptibility to clothianidin affects the efficacy of SumiShield® 50WG, one of the two newly approved formulations, which contains 50% clothianidin. We simultaneously monitored susceptibility to clothianidin and to SumiShield® 50WG, testing adults of *Anopheles gambiae*, *An. coluzzii* and *Culex* sp collected from urban, suburban and agricultural areas of Yaoundé. We found that the level of susceptibility to the active ingredient predicted the efficacy of SumiShield® 50WG. This formulation was very potent against populations that achieved 100% mortality within 72 h of exposure to a discriminating dose of clothianidin. By contrast, mortality leveled off at 75.4 ± 3.5 % within 7 days of exposure to SumiShield® 50WG in *An. gambiae* adults collected from a farm where spraying of acetamiprid and imidacloprid is driving cross-resistance to clothianidin. These findings indicate that more potent formulations of clothianidin or different insecticides should be prioritized in areas where resistance is emerging.

## Introduction

Malaria prevention in sub-Saharan Africa relies on two core interventions which used chemical insecticides ^1^. Indoor residual spraying (IRS) and long-lasting insecticidal nets (LLINs) have contributed to a substantial reduction in the disease burden over the last two decades, but the Achilles’ heal of both intervention measures is the development of insecticide resistance among vector populations ^2,3^. A limited number of active ingredients satisfy the criteria to be effectively deployed on a large scale in endemic areas ^4^. The spread of insecticide resistance poses a major challenge not only to the efficacy of IRS and LLINs, but also to their cost-effectiveness ^5^. Control programs have often attempted to mitigate the negative impacts of insecticide resistance by switching between active ingredients ^6^. For example, some endemic countries have progressively adopted more expensive alternatives such as carbamates and organophosphates for IRS to mitigate mosquito resistance to pyrethroids. This shift has been responsible for a decline in IRS coverage from 5% in 2010 to 2% in 2018 ^7^. Sequential deployment of insecticides is also associated with the development of multiple forms of phenotypic, genetic and behavioral resistance that quickly become established in wild mosquito populations ^1^. In order to reduce the likelihood of resistance development, the WHO’s global plan for insecticide resistance management in malaria vectors has recommended using active ingredients in rotations, combinations, mosaics and mixtures ^1,8^. However, for this management strategy to be successful, it is imperative to reinforce rotation programs with new chemicals which remain effective against mosquito populations that are currently tolerant to insecticides previously used in LLINs and IRS ^4^. In order to identify these alternatives, a number of agrochemicals have been tested and a few candidates have been selected for use in malaria vector control ^9–13^.

Clothianidin is manufactured into two new IRS formulations that have been evaluated and prequalified by the World Health Organization (WHO) ^4^. The two formulations are SumiShield® 50WG (50% clothianidin) developed by Sumitomo Chemical and Fludora^®^ Fusion (Bayer Environmental Science) which is a mixture of 500 g/kg clothianidin and 62.5 g/kg deltamethrin ^14,15^. There are currently 8 registered neonicotinoids, including clothianidin, which have become the most widely used agricultural pesticides worldwide ^16,17^. Neonicotinoids target the nicotinic acetylcholine receptor (nAChR) in the insect central nervous system and cause over-stimulation, which may result in paralysis and death ^17^. Although neonicotinoids are neurotoxic insecticides, their mode of action is sufficiently distinct, thus limiting the risk of cross-resistance with other neurotoxic chemicals such as pyrethroids.

Field trials of IRS formulations containing clothianidin have revealed long-lasting insecticidal activity against different vector species on diverse surfaces ^18–22^. Thus, clothianidin is considered a reliable alternative to control malaria vectors in areas with high pre-existing resistance to multiple insecticide classes. In preparation for rollout of clothianidin formulations as part of national IRS rotation strategies, the U.S. President’s Malaria Initiative has conducted susceptibility testing in anopheline populations from 16 African countries ^23^. This investigation suggested that most populations of the major vectors of *Plasmodium* parasites across the continent were susceptible to filter papers impregnated with SumiShield® 50WG at a diagnostic dose of 2% (w/v) clothianidin. While the findings of this study are encouraging, targeted testing of populations from agricultural regions where *Anopheles* mosquitoes are more likely to develop resistance to neonicotinoids should be conducted alongside continental-scale surveys.

The role of agricultural pesticides spraying in the development of resistance to insecticides used in malaria mosquito control has been widely documented ^24–31^. It has recently been demonstrated that *An. gambiae* larvae collected from farming areas where formulations of acetamiprid and imidacloprid, two neonicotinoids, are intensively used for crop protection can grow and emerge in water containing doses of neonicotinoids that were lethal to susceptible strains ^32^. This finding suggested that larval exposure to pesticide residues is selecting for cross-resistance to several neonicotinoids including clothianidin. Additionally, susceptibility tests conducted on adults confirmed resistance to thiamethoxam, imidacloprid and acetamiprid in some rural populations of *An. gambiae* ^33,34^. Neonicotinoids are highly soluble in water and can persist for months in aerobic soils, therefore making contamination of standing waters, which serve as mosquito breeding sites in farming areas very likely ^35–37^. Published information on the use of neonicotinoids across Sub-Saharan Africa is lacking, but preliminary reports from Cameroon, Tanzania and Ivory Coast suggested that hundreds of commercial formulations of neonicotinoids have been registered for use in crop protection ^38–41^.

Agriculture is one of the main activities in sub-Saharan Africa. The widespread application of neonicotinoids in crop protection could become a major challenge to the efficacy of repurposed chemicals such as clothianidin. Prior to rollout of formulations containing clothianidin, it is thus crucial to specifically test their efficacy against *Anopheles* populations that are showing signs of resistance to the active ingredient. Here we conducted intensive testing of *Anopheles* and *Culex* adult mosquito populations from five locations centered on one of the largest urban farms in Cameroon, using standard bioassays. Our aim was to determine if neonicotinoid tolerance observed in *An. gambiae* larvae and adults could affect the efficacy of SumiShield® 50WG. We found that as previously demonstrated by continent-wide surveys ^23^, SumiShield® 50WG is very effective against clothianidin-susceptible populations of *An. coluzzii* and *An. gambiae*. However, its efficacy is significantly reduced in *An. gambiae* adults displaying low mortality to clothianidin (150 µg/ml). These findings suggest that the effectiveness of clothianidin and its formulations should be monitored in depth in agricultural settings, and that alternative chemicals should be considered in areas where resistance to the active ingredient is emerging.

## Results

### 1. Evidence of clothianidin resistance in *Anopheles gambiae*

We used Centers for Disease Control (CDC) bottle bioassays to test a discriminating dose of clothianidin against a total of 1665 wild mosquitoes belonging to three species: *An. gambiae* (n = 912), *An. coluzzii* (n = 673) and *Culex* sp (n = 132) collected from 6 sampling sites. To validate our bioassay protocol, we first analyzed susceptibility in 554 individuals of two laboratory colonies: *An. gambiae* Kisumu (n = 326) and *An. coluzzii* Ngousso (n = 228). Results showed that the two laboratory strains were fully susceptible to clothianidin, reaching 100% mortality within 1-3 days of exposure (Fig. 2A). Among field mosquitoes, *Culex* sp. populations were the most susceptible to clothianidin, achieving 100% mortality approximately 24 h post exposure (Fig. 2A). A diagnostic PCR confirmed that specimens collected from the two urbanized sites were 100% *An. coluzzii*. These *An. coluzzii* populations were also susceptible to clothianidin, but 100% mortality was reached between the second and the fourth day. In *An. gambiae* by contrast, the overall mortality was only 58.2 ± 5.2 % in all 817 individuals tested from 4 locations, suggesting that some populations of this species have developed resistance to clothianidin. Knockdown at 60 min was generally low, except in field populations of *An. coluzzii* (81.8 ± 2.9%). Knockdown within 60 min of exposure ranged from 42 ± 11.0 % to 53.9 ± 9.9 % between lab strains and from 32.1 ± 5.0 % to 36.5 ± 12.2 % among field populations (Fig. 3A). Consistent with their reduced susceptibility to clothianidin, wild populations of *An. gambiae* displayed the lowest knockdown (32.1 ± 5.0 %).

**Figure 1:**
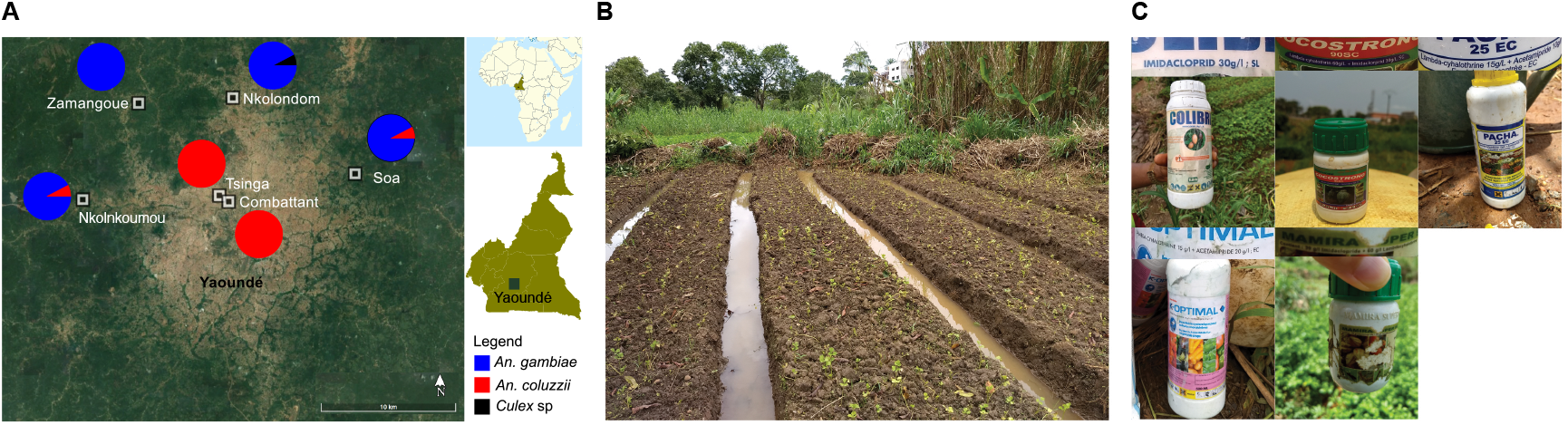
Description of the study site. (A) Map of the sampling locations where *Anopheles* and *Culex* mosquitoes were collected to evaluate susceptibility to clothianidin. (B) Picture of a larval breeding site in Nkolondom. (C) Examples of empty containers found in farms indicating the use of imidacloprid and acetamiprid formulations.

### 2. Resistance is stronger in populations from agricultural areas

To examine the link between larval exposure to agricultural neonicotinoids and the development of clothianidin resistance in *An. gambiae,* we compared the susceptibility profile of adult populations from different collection sites. Based on the diagnostic PCR, larvae collected from the farm or from one of the suburban sites (Zamengoué) were 100% *An. gambiae*. Samples from the two semi-rural sites, Soa and Nkolnkoumou, were ∼80% *An. gambiae*/20% *An. coluzzii*. All the four sites were treated as *An. gambiae* habitats. Mortality rates varied along a geographic gradient, ranging from susceptibility in Soa and Nkolnkoumou to resistance in samples originating from the agricultural site, Nkolondom. Only 46.5 ± 5.7 % of tested individuals from the agricultural site died between the first and the seventh day post-exposure (Fig. 2B). Susceptibility of adults from Nkolondom was significantly low compared with other *An. gambiae* sites (Wilcoxon Rank Sum test, p < 0.05). Knockdown at 60 min was also significantly lower (20.1 ± 4.1 %) in samples from the agricultural area (Wilcoxon Rank Sum test, p < 0.05) (Fig. 3B).

**Figure 2:**
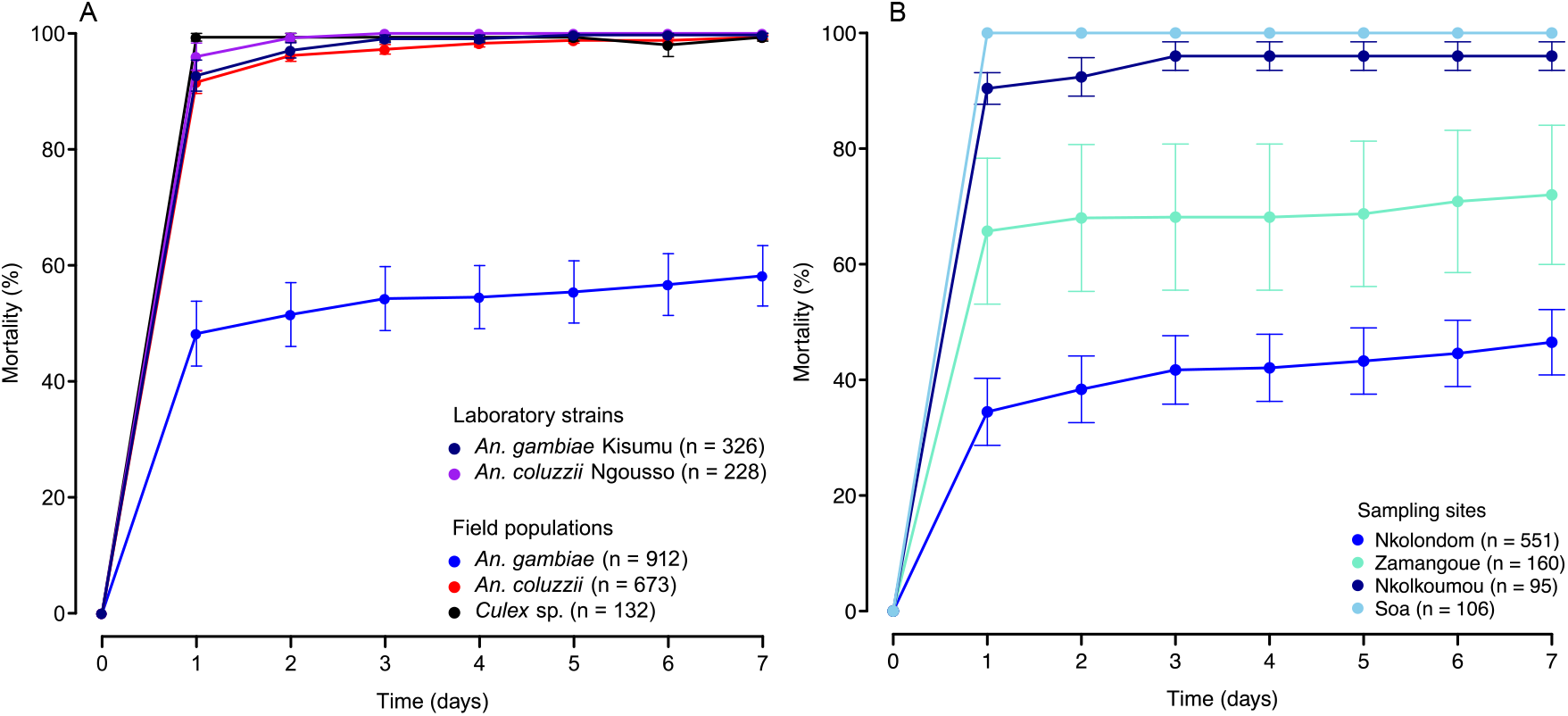
Baseline susceptibility of laboratory strains and field mosquito populations to clothianidin monitored for 7 days using CDC bottle bioassays. (A) Mortality values of *Anopheles* and *Culex* female adults exposed to 150 µg/ml clothianidin. (B) Gradient of susceptibility revealed in wild populations of *An. gambiae*. Error bars represent the standard error of the mean and (n) the number of individuals tested.

### 3. PBO is a synergist of clothianidin

Mortality observed in clothianidin-resistant populations of *An. gambiae* from the agricultural site increased from 46.5 ± 5.7 % without PBO to 92.7 ± 3.7 % when adult mosquitoes were pre-exposed to the synergist (Wilcoxon rank sum test, p=0.015) (Fig. 3C). This result suggested that metabolic detoxification involving cytochrome P450 enzymes contributes to the development of resistance to clothianidin in *An. gambiae*.

**Figure 3:**
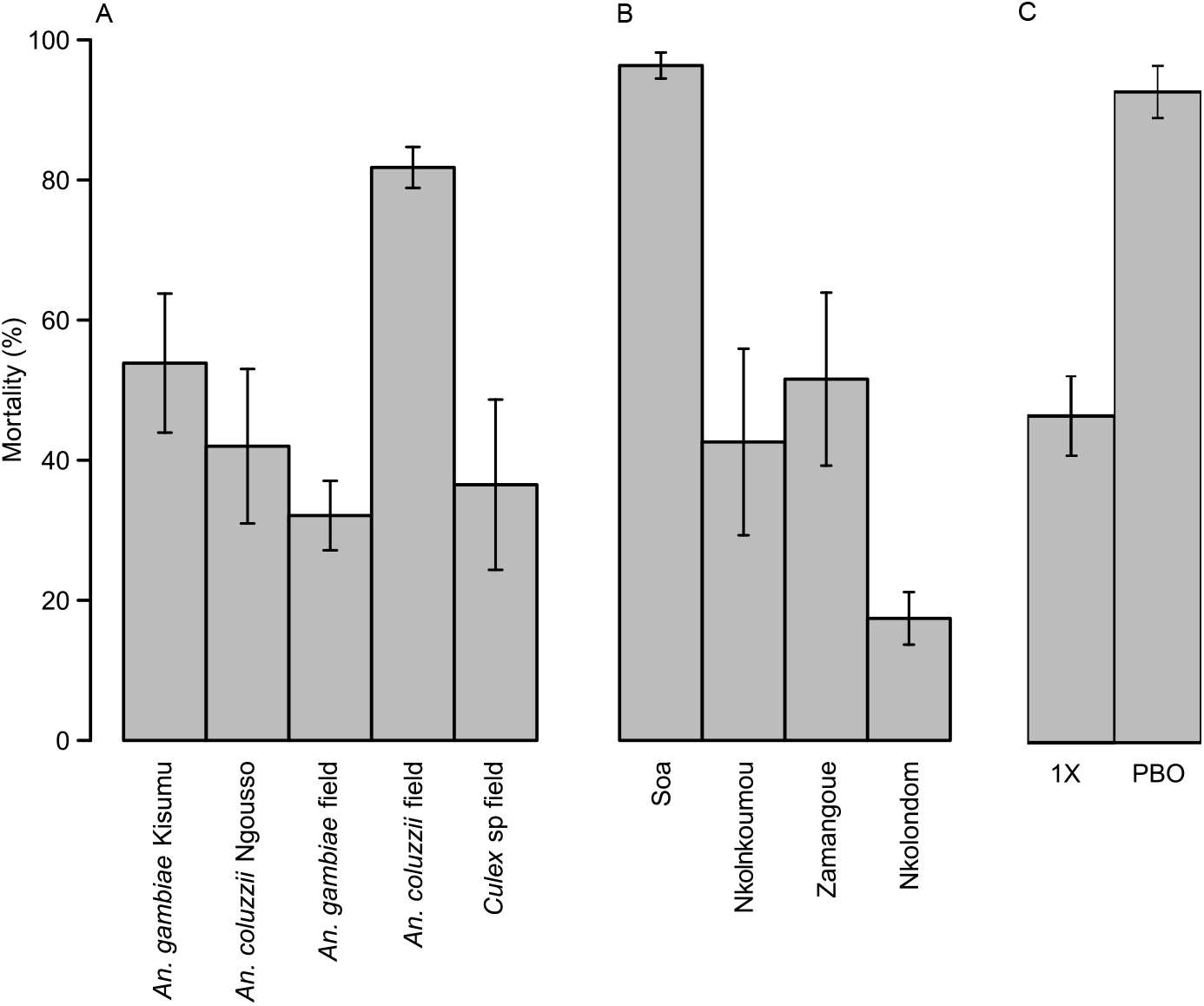
Knockdowns values and synergistic effect of PBO. Knockdowns after 1 h exposure to 150 µg/ml clothianidin were compared between *Anopheles* and *Culex* species (A) and between field populations of *An. gambiae* (B). Standard test (1X) and synergist bioassay (PBO) showing increase in mortality due to PBO (C). Standard errors of the mean are shown as vertical bars.

### 4. Resistance to clothianidin reduces the efficacy of SumiShield® 50WG

To determine to what extent clothianidin resistance could impact the efficacy of manufactured formulations, we used WHO tube tests to estimate the susceptibility of young female adult mosquitoes to SumiShield® 50WG. This formulation corresponds to ∼25-fold the discriminating dose of clothianidin used in CDC bottle assays. We tested one *An. coluzzii* population from urban areas (Combattant), one susceptible *An. gambiae* population (Nkolnkoumou) and resistant mosquitoes from Nkolondom. We used the laboratory strain *An. coluzzii* Ngousso as negative control. As expected, the lab strain was susceptible to SumiShield® 50WG, reaching 100% within 48 h of exposure to the formulation (Fig. 4). Among the field populations, susceptibility to clothianidin as revealed by bottle bioassays was a good predictor of the efficacy of SumiShield® 50WG. Both *An. coluzzii* and *An. gambiae* populations that were fully susceptible to the active ingredient were also susceptible to the IRS formulation. By contrast, *An. gambiae* populations from Nkolondom that were resistant to 150 µg/ml clothianidin in bottle bioassays were less susceptible to SumiShield® 50WG as well. Mortality rates against filter papers impregnated with SumiShield® 50WG at a diagnostic dose of 2% (w/v) clothianidin leveled off at 75.4 ± 3.5 % after 7 days of holding period. This finding suggests that *Anopheles* populations with low mortality to 150 µg/ml clothianidin could be challenging to control with current neonicotinoid formulations.

**Figure 4:**
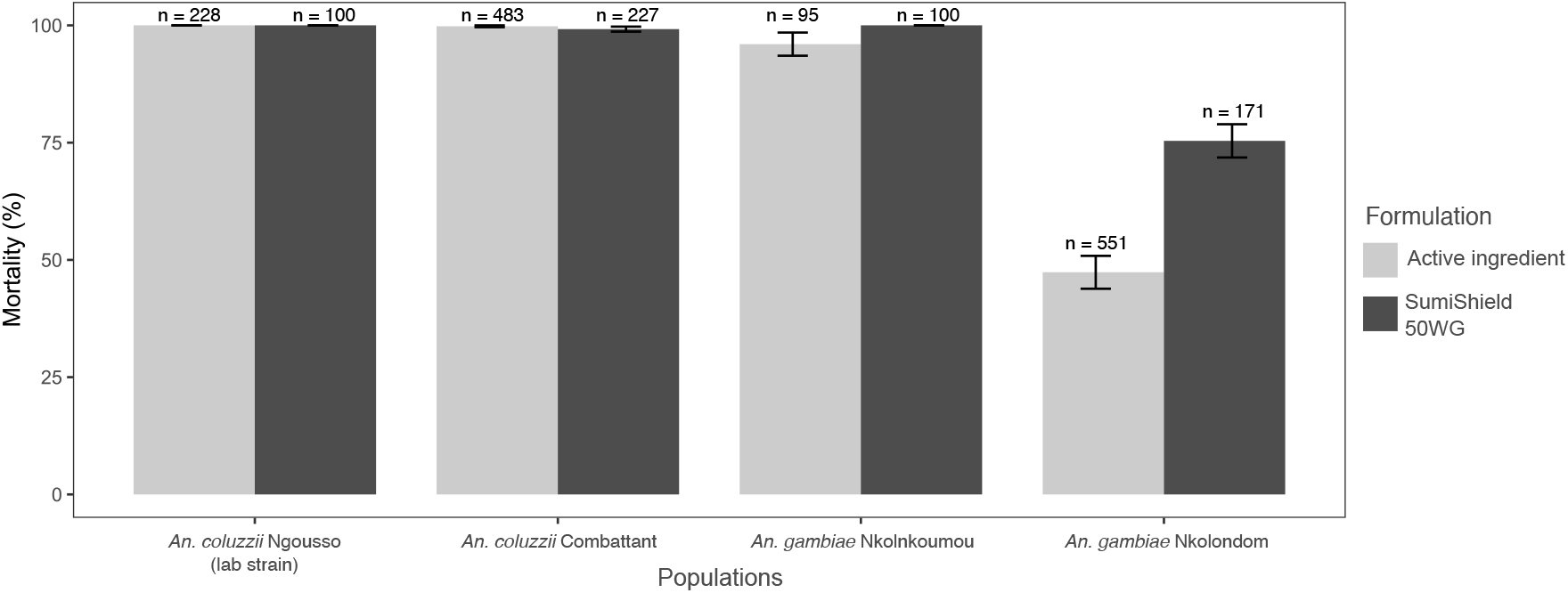
Relationship between susceptibility to clothianidin as revealed by CDC bottle bioassays and efficacy of SumiShield® 50WG evaluated with WHO tube tests.

## Discussion

Widespread pyrethroid resistance has been associated with a decline in efficacy of LLINs and IRS in several countries ^57–59^. Here we have provided evidence that the efficacy of some insecticides may have started to decline in some vector populations. The efficacy of SumiShield® 50WG, a formulation of clothianidin prequalified for IRS is reduced in *An. gambiae* mosquitoes that have evolved resistance to the active ingredient.

We first evaluated the susceptibility of *Anopheles* and *Culex* mosquitoes to clothianidin using CDC bottle bioassays. We used a protocol that differed slightly from the WHO standard operating procedure for testing the susceptibility of adult mosquitoes to clothianidin ^48,60^. We made the choice not to use a vegetable oil surfactant as suggested by the standard operating procedure because it has been demonstrated that these adjuvants are synergist of neonicotinoids ^34^. Additionally, at 150 µg/ml, the discriminating dose used, clothianidin was soluble in ethanol when the solution was allowed to rest for at least 24 h before use, and there was therefore no need to add a surfactant.

Using a discriminating dose of 150 µg/ml in CDC bottle bioassays, we successfully detected resistance to clothianidin in wild *Anopheles* mosquitoes. Knockdowns after 1 h exposure to the insecticide were low and had little discriminative power. Other studies assessing the susceptibility of adults *Anopheles* mosquitoes to clothianidin have reported low knockdowns, which could be due to the fact that clothianidin act slowly compared with other neurotoxic insecticides ^17,23,50,53^. Contrary to knockdowns, monitoring mortality rates for seven days provided a reliable assessment of susceptibility to clothianidin in adult mosquitoes. We first observed that susceptibility vary between species. *An. coluzzii* adults collected from urban areas of Yaoundé were susceptible to clothianidin. As crop cultivation associated with neonicotinoid spraying is less frequent in urban areas, *An. coluzzii* larvae from urbanized settings in Yaoundé are presumably less exposed to neonicotinoids residues and are therefore less likely to develop resistance to neonicotinoids. In *An. gambiae* however, the situation was more complex, with a gradient of susceptibility to clothianidin established among suburban and rural populations. Populations from a farm where neonicotinoids are used weekly for crop protection were the most resistant to clothianidin (Fig. 1C). Indeed, during our field survey in the Nkolondom farm, we collected empty containers of imidacloprid and acetamiprid confirming their use (Fig. 1C). Dozens of formulations of the two insecticides are freely sold in local stores in Yaoundé and are highly valued by farmers ^38,41^. The results presented in the current study are based on field surveys that were conducted between 2019 and 2020. The findings have been supported by monitoring that continues from 2020 to 2022 and confirmed patterns of susceptibility to clothianidin observed in precedent years in *Anopheles* mosquitoes from Yaoundé and its neighboring rural areas ^33^. These surveys have combined larval tests and adult bioassays to reveal that neonicotinoid resistance is emerging in *An. gambiae* populations from the equatorial forest region of Cameroon, especially in areas where larvae are chronically exposed to pesticide residues ^32–34^. There is ample evidence that *An. gambiae* larvae and adults from several villages around the city of Yaoundé are currently resistant to imidacloprid, acetamiprid and thiamethoxam, three neonicotinoids that are among the most widely used crop protection chemicals in Cameroon ^32,33,38,41^. A study conducted in Ivory Coast has also observed resistance to imidacloprid and acetamiprid in *An. coluzzii* correlated with agricultural activities ^61^.

Intriguingly, adults of *Culex sp* whose larvae were collected from the same breeding sites as *An. gambiae* in Nkolondom were fully susceptible to clothianidin. However, it is well known aquatic invertebrates have variable responses and threshold of susceptibility to the lethal and sublethal effects caused by neonicotinoid contaminants^62^. Although *Culex* sp populations were more directly impacted by the lethal toxicity of clothianidin, they likely have developed other physiological and/or behavioral adjustments enabling them to adapt to neonicotinoid residues in farms ^63,64^.

Bioassay tests using CDC bottles coated with the synergist PBO prior to exposure to clothianidin showed a drastic increase in mortality in resistant populations. This suggested that Cytochrome P450 enzymes (CYPs) play a primarily role in neonicotinoid resistance in *An. gambiae*. Acetamiprid resistance in *An. gambiae* is also highly dependent on metabolic detoxification mediated by CYPs ^33^. More generally, neonicotinoid resistance in wild populations of many crop pests, primarily those of the order *Hemiptera* (aphids, whiteflies, and planthoppers), is associated with overexpression of one or several CYP enzymes ^65–68^.

The spread of pyrethroid resistance has caused a decline in the effectiveness of LIINs and IRS ^57,58^. There is a risk that resistance will also undermine the efficacy of new products such as SumiShield®® 50WG and Fludora® Fusion. Precisely, larval bioassays showed that the intensity of resistance to clothianidin in *An. gambiae* collected from Nkolondom is currently similar to that of deltamethrin ^32^. Therefore, even without any large-scale deployment of clothianidin in vector control, its efficacy may already be as reduced as that of pyrethroids in some populations ^69^. In our study, we have revealed that the efficacy of SumiShield®® 50WG is declining in clothianidin-resistant populations. We did not test Fludora® Fusion, and it remains to be determined if the dual action of clothianidin and deltamethrin will result in higher efficacy against resistant populations. However, a recent experiment has demonstrated that exposure of *An. gambiae* larvae to sublethal doses of a mixture of different types of agrochemicals increased the tolerance of adults to both clothianidin and Fludora® Fusion ^70^. This suggested that any formulation is at risk of losing its efficacy if clothianidin resistance spreads among anopheline populations.

## Conclusions

The current study shows that the new insecticide clothianidin may have reduced efficacy in some agricultural regions of Sub-Saharan Africa due to pre-existing levels of resistance among mosquito populations. These findings suggest that prior to inclusion of agrochemicals in resistance management programs, their efficacy should be particularly scrutinized in areas where pesticides of the same class are being used for agricultural pest management.

## Methods

### Study sites

We collected mosquitoes from four suburban sites in the outskirts of Yaoundé, the capital of Cameroon (Figure 1A). We also tested samples collected from two densely urbanized areas of the city. Approval to conduct a study in the Center region (N°: 1-140/L/MINSANTE/SG/RDPH-Ce), ethical clearance (N°: 1-141/CRERSH/2020) and research permit (N°: 000133/MINRESI/B00/C00/C10/C13) were granted by the ministry of public health and the ministry of scientific research and innovation of Cameroon. The suburban sites included Nkolondom (3°56’43” N, 11°31’01” E), situated approximately 7 km west of Yaoundé. Since 1985, a swampy area in Nkolondom is exploited for intensive crop cultivation ^42^. In 2020, the farm which attracted at least 100 workers was subdivided into 100-200 mosaics of ∼20m^2^ adjacent plots dedicated to the cultivation of aromatic herbs, amaranth and lettuce (Fig. 1B). Standing water between ridges and furrows provide mosquito breeding sites, which maintain large larval populations of *An. gambiae* and *Culex sp* throughout the year ^32–34^.

### Mosquito populations

We sampled and cumulatively tested mosquitoes from the six above-mentioned sites between 2019 and 2020. *An. gambiae sensu lato* larvae were collected from standing water using dippers and transported in plastic containers to the insectary where they were reared to adults under standard laboratory conditions of 25–27°C, 70–90% relative humidity and a 12:12 h light/dark photoperiod ^43^. The two sibling species *An. gambiae sensu stricto* (hereafter *An. gambiae*) and *An. coluzzii* are the dominant malaria vectors in Yaoundé ^44,45^. *An. coluzzii* is found exclusively in the most urbanized neighborhoods while *An. gambiae* is the only species present in neighboring rural areas. To determine which species between *An. gambiae* and *An. coluzzii* was collected from each site, we genotyped 50 mosquito samples using a diagnostic PCR ^46^. *Culex* sp larvae that occur in the same breeding sites as *An. gambiae* in the Nkolondom farm were also sampled, reared to adults and tested.

### Clothianidin susceptibility testing

The susceptibility of adult mosquitoes that emerged from field-collected larvae was tested against clothianidin using CDC bottle assays ^47^. The bioassay procedure followed a modified version of the WHO standard operating procedure for testing the susceptibility of adult mosquitoes to clothianidin ^48,49^. Precisely, we did not used the vegetable oil surfactant, Mero® (1% v/v), as adjuvant. A recent study showed that some vegetable oil surfactants are synergists of neonicotinoids, which when used in susceptibility testing lead to an overestimation of the insecticidal activity of the active ingredient ^34^. Instead, mortality was evaluated against the active ingredient alone dissolved in ethanol using a discriminating dose (i.e., the lowest dose at which 100% of adults from a susceptible population die) of 150 µg/ml as determined by a previous study ^50^. We prepared stock solutions using a technical-grade formulation of clothianidin (PESTANAL^®^, analytical standard, Sigma-Aldrich, Dorset, United Kingdom) and absolute ethanol as solvent. The solutions were conserved at 4°C in the dark for at least 24 h before use to maximize solubility of clothianidin. Each Wheaton 250-ml bottle and its cap were coated with 1 ml of a solution containing 150 µg/ml clothianidin in ethanol following the CDC guidelines ^47^. For each bioassay, we used four test bottles coated with clothianidin and two control bottles coated with 1 ml of absolute ethanol. All bottles were wrapped in aluminum foil and allowed to dry for 24 h for complete evaporation of solvent before use. Coated bottles were not reused and were washed three times in warm soapy water and allowed to dry for 24 h between experiments. 20 to 25 3-5-day-old females were aspired from mosquito cages and released into one of the test bottles where they were exposed to the active ingredient or into control bottles containing ethanol for 1 h. After the exposure period, mosquitoes were transferred into a paper cup and provided with 10% sugar solution. Knockdowns were scored immediately and mortality was monitored every 24 h for seven consecutive days. We used adults from two susceptible strains as controls: *An. gambiae* Kisumu and *An. coluzzii* Ngousso. Both strains are known to be susceptible to pyrethroid, carbamate, organochlorine and organophosphate insecticides.

### Synergist bioassay

We used CDC bottle assays to test if the synergist piperonyl butoxide (PBO) could enhance the potency of clothianidin. PBO is an inhibitor of oxidases and non-specific esterases ^51^. A solution of 400 µg/ml PBO was prepared by diluting PBO with absolute ethanol. The solution was mixed and stored in the dark at 4°C before use. Wheaton bottles and their cap were coated with 1 ml of the PBO solution, wrapped in aluminum foil and allowed to dry for 24 h before the test. 100 to 150 female mosquitoes aged between 3 and 5 days were pre-exposed for 1 h to PBO-coated bottles or to control bottles coated with ethanol. Mosquitoes were then removed from the bottles and batches of 20 to 25 individuals were introduced into new bottles coated with 150 µg/ml clothianidin or with ethanol. After 1 h, mosquitoes were transferred into a paper cup and provided with 10% sugar solution. Knockdowns were scored and mortality was monitored every 24 h for seven consecutive days in standard laboratory conditions. We compared mortality rates with or without the synergist to determine if oxydase inhibition by PBO affected the level of susceptibility.

### Efficacy of SumiShield® 50WG

We used WHO tube tests to assess the efficacy of SumiShield® 50WG against adult mosquitoes displaying a gradient of tolerance to clothianidin ^52^. We used a discriminating dose of 2% w/v clothianidin (13.2 mg active ingredient per paper) following the manufacturer’s recommendation ^23,53^. We prepared a stock solution by diluting 264 mg SumiShield® 50WG in 20 ml distilled water. We impregnated Whatman filter papers (12 x 15 cm) containing 13.2 mg clothianidin each using 2 ml of the insecticide solution as described in ^54^. Control filter papers were impregnated using 2 ml of distilled water. Treated filter papers were allowed to dry overnight and were kept in foil at 4°C until use. To carry out bioassay tests, we aspired 20 to 25 3-5-day-old female mosquitoes from mosquito cages that we introduced into each of the test tubes containing clothianidin-impregnated papers. We concomitantly released 20-25 mosquitoes into each control tube. After 1 h, mosquitoes were transferred into holding tubes and provided with 10% sugar solution, and knockdown time was recorded. As with CDC bottle tests mortality was scored every day until day 7.

### Data analysis

We used mortality rates to evaluate the efficacy of clothianidin against laboratory and field mosquitoes. All tests with mortality > 20% at day 7 in controls were discarded. We used Abbott’s formula to correct mortality rates of tests when 5–20% individuals died between day 1 and day 7 in the corresponding controls ^55^. Following the WHO guidelines on insecticide susceptibility, mosquito populations were considered susceptible if mortality at day 7 was ≥ 98% and resistant if mortality was less than 90%. Mortality rates between 90% and 97% indicate that the presence of resistant genes in the vector population must be confirmed by additional tests ^52^. We used a Wilcoxon rank sum test to determine if mean mortality rates were significantly different among tested populations. We performed all analyses using the R software V 4.2.2 ^56^.

## Author Contributions

CF: Conceptualization, Formal analysis, Investigation, Methodology, Writing – original draft; FA: Investigation, Methodology; MA: Investigation, Methodology; WT: Investigation, Methodology; CW: Resources, Supervision. CK: Conceptualization, Formal analysis, Funding acquisition, Investigation, Project administration, Writing – review & editing.

## Data Availability Statement

The data for this study have been presented within this article.

